# Pathogenic *BRCA1* mutations disrupt allosteric control by *BARD1*

**DOI:** 10.1101/2025.06.07.658438

**Authors:** Ayan Bhattacharjee, Gregory R. Bowman

**Affiliations:** Departments of Biochemistry & Biophysics and Bioengineering, University of Pennsylvania, Philadelphia, PA 19104, United States

## Abstract

Mechanistic insight into biophysical perturbations caused by pathogenic missense mutations is highly valuable information for the rational design of therapeutics. For hereditary breast and ovarian cancer, multiple pathogenic mutations in the N-terminal domain of *BRCA1* have been reported in patients. How exactly these mutations disrupt the catalytic activity of *BRCA1*, and thereby lead to oncogenesis, is unknown. Here, we posit that the mechanism of pathogenesis is tied to how binding of *BARD1* activates *BRCA1* for E3 ligase activity. We use atomistic molecular dynamics simulations and Markov state modeling to uncover how *BARD1* selects for active conformational states of *BRCA1*. We show that the helix bundle, where *BARD1* binds, is allosterically coupled to the E2 interface. Furthermore, we show that *BARD1* selects for conformational states that are pre-organized for E3 activity. Lastly, we show that pathogenic mutations allosterically destabilize active states, whereas hyperactive mutations constitutively increase their likelihood. These results provide a concrete strategy supported by mechanistic insight for the design of restorative small molecules targeting *BRCA1*.

## Introduction

Understanding the mechanisms by which mutations modulate protein function is a grand challenge with significant implications for our ability to treat diseases and predict the implications of variants of unknown significance^1^. Missense mutations in proteins are a common source of pathogenesis for many diseases^2^, including cancers. In cancer genomes, missense mutations are the most common type of protein-coding mutation^3^. Significant progress has been made in high-throughput prediction of oncogenic missense mutations^4,5^, but the mechanisms by which missense mutations cause gain or loss-of-function are often unclear without structural insights^6^. Thorough understanding of these mechanisms, however, have formed the foundation of therapeutic strategies for many cancers that have been successful in the clinic^7^. Unraveling how missense mutations cause disease phenotypes at the molecular level leads to actionable strategies for rational drug discovery.

Three decades of research have established breast cancer type 1 susceptibility protein (*BRCA1*)^8–11^ as a tumor suppressor for breast and ovarian cancer, one of the most common cancers globally^12^. Missense mutations comprise 36% of all *BRCA1* mutations^13^. Many of these mutations are found in the N-terminal domain of *BRCA1*, which is a RING-type^14^ (“really interesting new gene”) E3 ubiquitin ligase^15,16^. Formation of a heterodimer between *BRCA1* and *BARD1*(breast cancer associated RING domain 1) is required for *BRCA1* to be catalytically active and facilitate ubiquitin transfer as an E3 ligase^17–19^. The N-terminal domains of *BRCA1* and *BARD1* are RING domains, each containing two zinc fingers^20^. Zinc finger II of *BRCA1* forms part of the interface to partner E2 ligases^21^. This heterodimer is formed by interaction between the N-terminal domains of *BRCA1* and *BARD1* with an extensive hydrophobic interface formed by complementary helix bundles (Fig 1a-b). The *BRCA1/BARD1* heterodimer has many ubiquitination substrates^22^, such as the recently identified complex with histone 2A in nucleosomes^23–26^.

**Figure 1.**
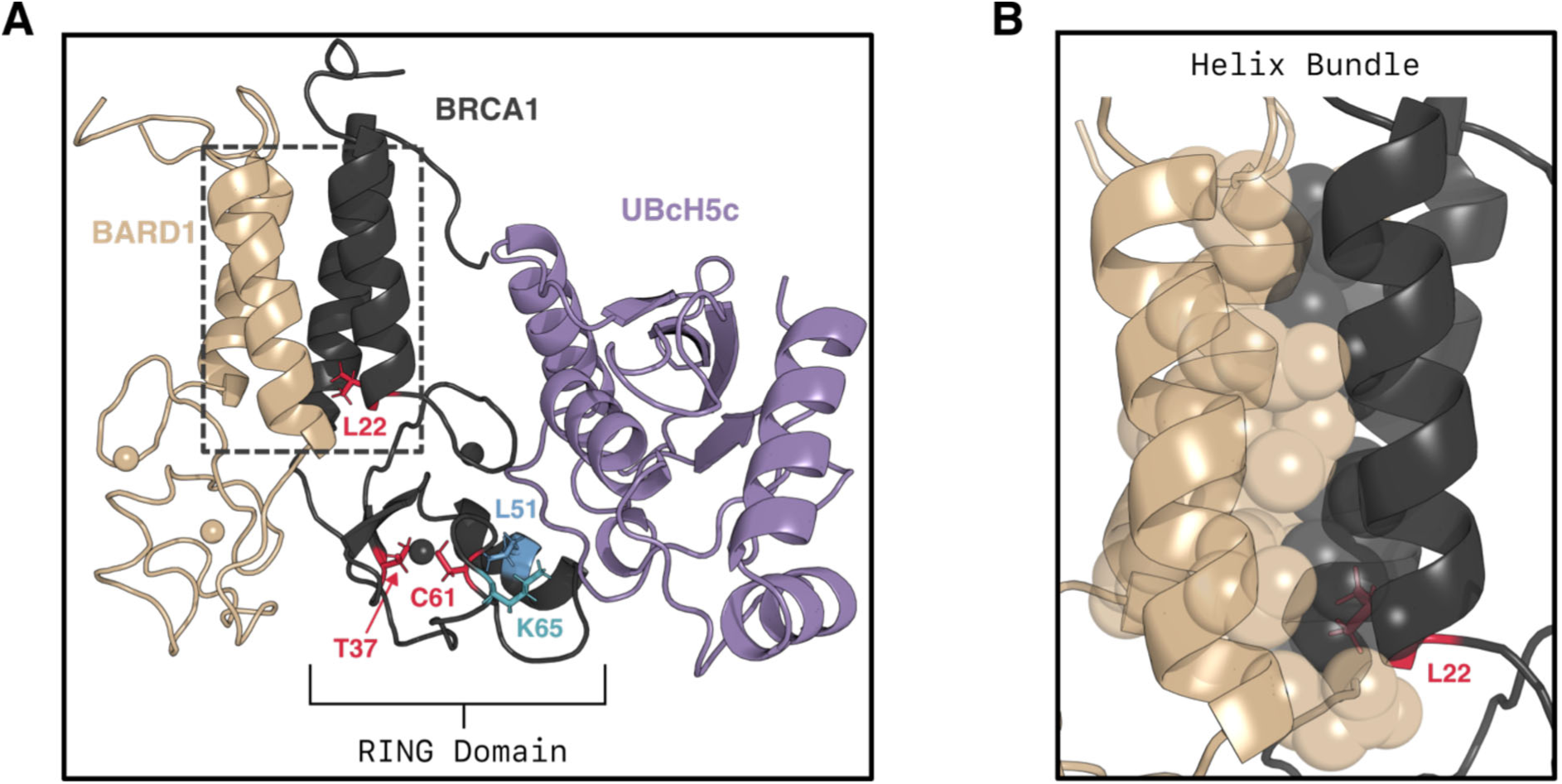
Structure of the BRCA1 RING domain bound to BARD1 and the E2 ligase UBcH5c (PDB: 7LYB). A) Ternary structure highlighting the locations of three pathogenic mutations in red sticks (L22S, T37K, and C61G) and mutations in a hyperactive double mutant in blue sticks (L51W) and cyan sticks (K65). K65 is the canonical linchpin residue in RING-type E3 ligases. The dashed box corresponds to the zoomed inset in panel B. B) Zoom in on the helix bundle at the BRCA1/BARD1 interface. Residues in BRCA1 and BARD1 making key hydrophobic contacts with each other are shown in translucent spheres.

The precise mechanism for how *BARD1*-binding activates *BRCA1* and how pathogenic mutations disrupt activity is unknown. Notably, the *BARD1* RING does not interact with E2 ligases such as *UBcH5c* despite being necessary for catalytic activity^27^. It has been proposed that *BARD1*-binding selects for active state conformations of *BRCA1*, but never confirmed experimentally as NMR studies of *BRCA1* RING monomer were not amenable^20^. Furthermore, pathogenic variants are known to disrupt catalytic activity in the RING domain, but the mechanism is unclear. Mutations to metal-coordinating residues in zinc finger II (e.g. C61G) result in loss-of-function and local structural instability but do not abrogate *BARD1* binding^27^, while the biophysical effects of others (e.g. T37K) have not been characterized at all. Reasoning that a general structural mechanism of pathogenesis would involve disruption of *BARD1*-mediated activation of *BRCA1*, we sought to jointly answer these questions.

Given the necessity of *BARD1*-binding for *BRCA1* E3 ligase activity, we hypothesized that there would be a direct allosteric relationship between the *BARD1*-interface and the *E2*-interface in the RING domain of *BRCA1*. The presence of allostery is detectable in the structural ensemble of a protein if a conditional population shift is observed^28^. Recently, all-atom molecular dynamics simulations have emerged as a leading method to determine the ensemble of complex biomolecular systems^1,29^, and thereby the presence of allostery^30,31^.

Hence, to assess our hypothesis, we utilized atomistic molecular dynamics (MD) simulations and Markov state models (MSMs)^32^ to elucidate the structural ensembles of *BRCA1,* the *BRCA1*/*BARD1* complex, and mutations known to increase or decrease activity.

## Results

### Isolated *BRCA1* rarely adopts functional conformations

Given that the *RING* domain of *BRCA1* is only catalytically active when bound to *BARD1*, we hypothesized that *BARD1* may stabilize active conformations of *BRCA1* that are otherwise rare. Prior experimental work provides indirect support for such a mechanism. NMR experiments have shown that *BARD1* does *not* form any contacts with E2 ligases as measured by chemical shift perturbations^20^. Additionally, Stewart et al. found that a “*RING*-less” variant of *BARD1* containing only the helix bundle activates *BRCA1* more than wild-type *BARD1*. Hence, we predicted that the interaction between the helical bundles of *BRCA1* and *BARD1* (Fig. 1b) would drive this conformational selection.

To explore this hypothesis, we used atomistic molecular dynamics simulations to map out the ensembles of structures that isolated *BRCA1* and the *BRCA1*/*BARD1* dimer adopt. Powered by the citizen scientists of Folding@home^33^, we conducted an aggregate ∼450 µs of simulations for both systems in explicit solvent. Separate MSMs were constructed for the *BRCA1* conformational ensembles in the monomer and heterodimer states using the software deeptime^34^, as described in the Materials & Methods. We then characterized how variable the ensembles were by computing the distribution of root-mean-square-deviations (RMSD’s) to the experimentally resolved *BRCA1/BARD1* structure. We then compared these distributions to identify direct evidence for conformational selection into active states by *BARD1*. If the isolated *BRCA1* ensemble differs significantly, this indicates that monomeric *BRCA1* often adopts conformation incompatible with binding *BRCA1*; alternatively, if both distributions are similar, that indicates an alternative mechanism is possible.

We find that indeed *BARD1* stabilizes *BRCA1* in a small subset of the broad ensemble of structures that isolated *BRCA1* can adopt. In simulation, monomeric *BRCA1* deviates more than heterodimeric *BRCA1* from the NMR structure determined by Brzovic et al. The distribution of root-mean-square deviation (RMSD) to the reference for the monomer has a higher mean and variance than the dimer (Fig 2a). We then verified that these deviations were not due to unfolding of the protein in simulation. For each residue in the protein we computed the probability that each residue exhibited secondary structure, assessed by the DSSP algorithm^35^, for both ensembles. Comparing these probabilities, we found that monomeric *BRCA1* mostly retains the same fold as heterodimeric *BRCA1* (Fig 2b), verifying that no unfolding polluted the RMSD measurement. Together, these data suggest that monomeric *BRCA1* rarely adopts conformations similar to that of the experimental structure of *BRCA1* in complex with *BARD1*.

**Figure 2.**
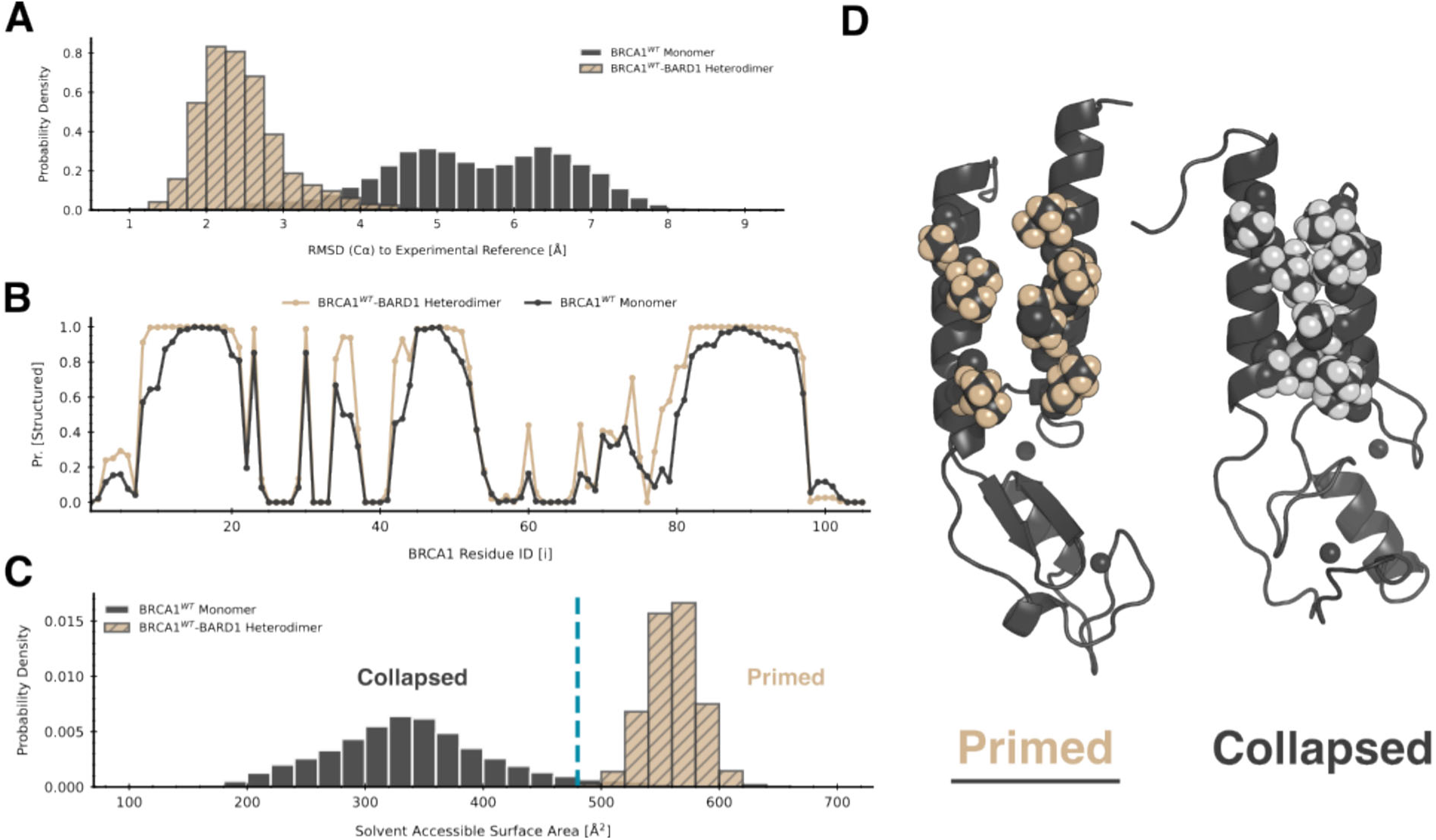
Monomeric BRCA1 only rarely adopts BARD1-binding competent conformations, instead tending to undergo hydrophobic collapse at the helix bundle, which causes a conformational change in the RING domain. A) The RMSD distribution of BRCA1 structures in simulations of the monomer and the heterodimer with respect to an experimentally determined reference (PDB: 7LYB). The spread of the monomer distribution (black) is wider than the heterodimer distribution (gold with black hatch marks), indicating the monomer has increased conformational heterogeneity. B) The probability of each residue being structured in the BRCA1 ensemble. A residue is considered structured if it is part of an alpha helix or beta sheet. Secondary structure elements are mostly conserved between heterodimer (gold) and monomer (black) simulations. C) The distribution of the exposed solvent accessible surface area (SASA) of the helix bundle. For heterodimer simulations (gold with black hatch marks), SASA is computed without BARD1 present; hence, a larger SASA indicates a larger binding interface. The monomer (black) prefers collapsed states with decreased SASA. The vertical blue line marks a cutoff between what we call primed and collapsed states. D) Representative structures of collapsed (right) and primed (left) states from simulations of the BRCA1 monomer. Residues necessary for making key hydrophobic contacts with BARD1 are shown in spheres. Collapsed states feature an inward motion of the helices, which pack the hydrophobic residues in each helix against each other.

To understand the structural features that distinguish conformations of *BRCA1* only seen in the absence of *BARD1* from those in the heterodimer, we further investigated the dynamics of the helix bundle. In particular, we wanted to assess the *BARD1*-binding compatibility of the interface in the two *BRCA1* ensembles. To quantify this, we compared the distribution of the helix bundle’s solvent accessible surface area (SASA) between monomeric and heterodimeric *BRCA1*. This was as computed as the sum of atomwise SASA’s for non-backbone atoms for the seven hydrophobic residues at the protein-protein interface (Fig. 1b) using enspara^36^. For simulations of the heterodimer, the calculation was done with *BARD1* absent. Therefore, a higher SASA indicates a larger interface for *BARD1* to bind where key sidechains are “facing out”, whereas a lower SASA indicates more intra-helix hydrophobic contacts that are suboptimal for *BARD1*-binding.

Consistent with our hypothesis that monomeric *BRCA1* often adopts conformations incompatible with binding *BARD1*, we found that the helix bundle SASA distribution of monomeric *BRCA1* has little overlap with that of heterodimeric *BRCA1* (Fig. 2c). Furthermore, the monomeric *BRCA1* ensemble exhibits less helix bundle SASA than heterodimer *BRCA1*. After visual inspection of conformational states at the left tail of the monomer distribution, we observed that monomeric *BRCA1* often inhabited states where the individual helices in the bundle would twist to bury each other (Fig. 2d), which we term collapsed. Conversely, at the right tail of the monomer distribution we observed conformations where the helix bundle was oriented such that the hydrophobic sidechains could interlock with *BARD1*, which we term primed states. Hence, we find that *BRCA1* strongly prefers collapsed states which are not compatible with *BARD1*-binding and rarely adopts primed states, which are more dimer-like and therefore functional.

### Allostery in *BRCA1* provides a means for *BARD1* to control *BRCA1*-E2 binding

We reasoned that the conformational differences described above could provide a means for *BARD1* to allosterically control whether *BRCA1* can bind partner E2 ligases such as UBcH5c^20,22,23,26^. It is known that that *BRCA1* is only active as an E3 ligase in the presence of *BARD1*^20^, but it remains unclear why. Cryo-EM structures of *BRCA1/BARD1* in complex with an E2 ligase and nucleosome substrates are very similar structurally to the isolated heteodimer, offering no particular explanation. We hypothesized that *BRCA1* conformations stabilized by *BARD1* are more competent to bind E2 ligases, and that there exists allosteric communication between the helix bundle and the *RING* domain to control these conformational changes.

To identify such allostery, we trained a DiffNet^37^ neural network to learn how *BRCA1*’s E2 interface was affected by the *BARD1*-binding interface (see *Methods*). DiffNets identify structural features that differentiate conformational ensembles by requiring that they classify a biochemical property of interest, encoded by a continuous label on a scale from 0 to 1. In this case, a natural choice is to have the label reflect E3 ligase activity. To achieve this, we initially gave all *BRCA1* structures from the heterodimer simulations a label of 1 and all *BRCA1* structures from the monomer simulations a label of 0. Then, we let DiffNets learn a representation of the protein with self-consistent labels. That is, similar structures from the monomer and heterodimer simulations must have the same label, and this label should be closer to 1 if the structure is more probable in the heterodimer simulations representing an active state and the label should be closer to 0 if it is more probable in simulations of the monomer that we presume to be inactive. Finally, we identify inter-residue distances that are correlated with the label to find structural features that distinguish inactive and active conformations.

As expected, we find that when *BRCA1* adopts conformations that are primed to bind *BARD1*, it also appears to be pre-organized to bind an E2 ligase. We find that the distance between I14 and K88 is correlated with the labels predicted by DiffNets, while the distance between S36 and L63 is anti-correlated with the DiffNets labels. Projecting the free energy landscape of *BRCA1* onto these two distances shows that they are anti-correlated (Figure 3A). Mapping these distances onto representative structures helps provide mechanistic insight (Figure 3B). Monomeric (inactive) *BRCA1* prefers structures where the I14-K88 distance is small, suggesting the helices are collapsed and incompetent to bind *BARD1*. At the same time, the S36-L63 distance is large, indicating disordering of the foot of *BRCA1* that includes the E2 ligase-binding site. In contrast, active *BRCA1* (bound to *BARD1*) favors conformations where the I14-K88 distance is large, consistent with the primed state of the helices that is competent to bind *BARD1*. Concomitantly, the S36-L63 distance is small, indicating proper folding of the foot of *BRCA1* such that it is competent to bind the E2 ligase. Hence, a *BARD1-*binding compatible helix bundle geometry is allosterically coupled to an E2-binding compatible local geometry in the *RING* domain.

**Figure 3.**
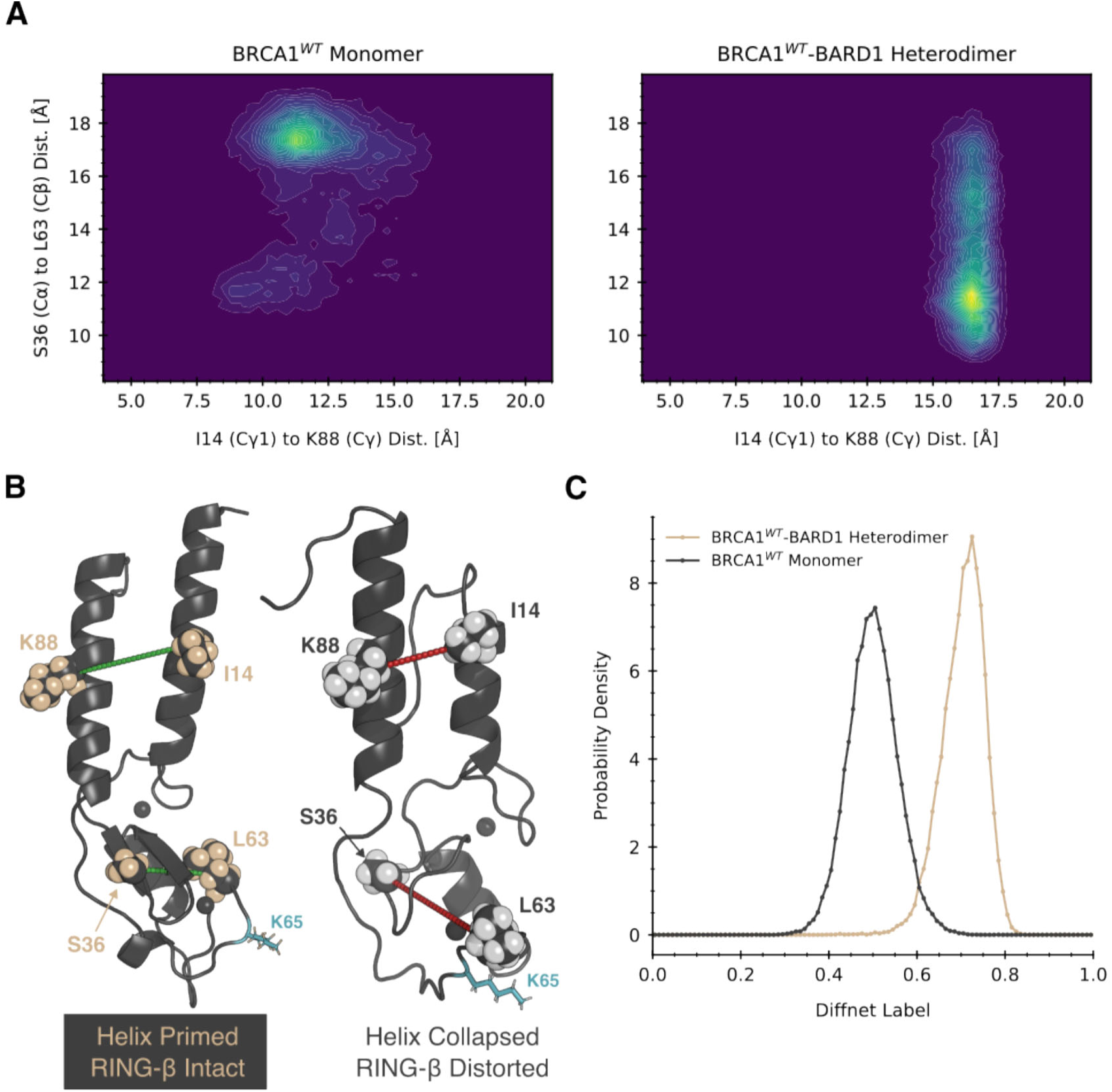
The structure of the BARD1-binding interface is allosterically coupled to the structure of the E2-binding interface. A) A projection of the BRCA1 energy landscape onto two interatomic distances which correlate with the DiffNets output label. The x-axis corresponds to the distance between the gamma carbons of I14 and K88, and the y-axis corresponds to the distance between the alpha carbon of S36 to the beta carbon of L63. A negative correlation is observed between these two distances. Comparing the two projections shows that monomeric BRCA1 rarely adopts BARD1-binding competent conformations. B) The structural depiction of the two distances in landscape in part (a). The distance of I14 to K88 reports on the state of hydrophobic collapse in the helix bundle, whereas S36 to L63 correlates to a reorientation of the second-zinc finger, a conformational change in the E2-binding cleft, and a disruption of the beta sheet in the RING finger. C) The DiffNet label distributions for BRCA1 structures from monomeric and hetero-dimeric simulations. DiffNet labels can represent any biochemical property of interest which is a function of conformation; here, zero represents inactive states and one represents active states.

### Pathogenic missense mutations allosterically disrupt linchpin priming

We hypothesized that the mechanism by which missense mutations reduce E3 ligase activity in BRCA1 is by preventing the conformational selection of catalytically active states. Three missense mutations reported in individuals which are labeled pathogenic in ClinVar^38^ were studied: L22S, T37K, and C61G. The mutation C61G is known to disrupt zinc-binding and cause local instability at the second zinc finger^27^. While residue L22 is located at the base of the helix bundle, it is not as exposed to *BARD1* relative to other hydrophobic residues at the interface (eg. L82). Residue T37 is located in the *RING* β-sheet, and it is unknown why mutation to K results in loss of E3 activity.

Because none of these mutations are located in the E2-binding interface (Fig. 1a), we reasoned that they would allosterically prevent the interface from being pre-organized for E3 ligase activity. Specifically, we hypothesized that pathogenic mutations would disrupt the optimal positioning of the “linchpin” residue, a common feature shared across *RING*-type ligases^39,40^ that is critical for ubiquitination and neddylation. The linchpin residue stabilizes the “closed” state of the E2∼Ubiquitin (Ub) conjugate, priming it for Ub transfer, by engaging in a hydrogen-bonding network with both E2 and Ub. Previous work has highlighted that residue K65 is the linchpin residue in *BRCA1*^41^; substitution of K65 to A results in a ligase-dead mutant, whereas mutation to R results in a catalytically hyperactive mutant.

To explore this hypothesis, we assessed the probability that the K65 linchpin is correctly positioned, or primed, for E3 ligase activity when *BRCA1* is activated by *BARD1* to an primed state for monomeric wild-type and mutants. We conducted an additional ∼350 µs of simulations for the three mutant systems. We analyzed priming of the K65 linchpin by comparing ensembles from simulations to a constructed structural model of the active *BRCA1*/*BARD1*/*UBcH5c*/Ubiquitin complex The model was constructed by alignment of the *BRCA1*/*BARD1* NMR model^19^ to the x-ray structure of the heterotrimeric complex *RNF4*/*UBcH5a*/Ubiquitin^42^. As expected, this model buries the sidechain of K65 residue into the E2/Ub interface, where the terminal amine forms hydrogen bonds with Q92 of UbcH5c and R72 of Ubiquitin (Fig. 4b). For each ensemble, the component structures were aligned to the *RING* domain of this model; then, the RMSD of the linchpin K65 in each structure was computed to the reference.

**Figure 4.**
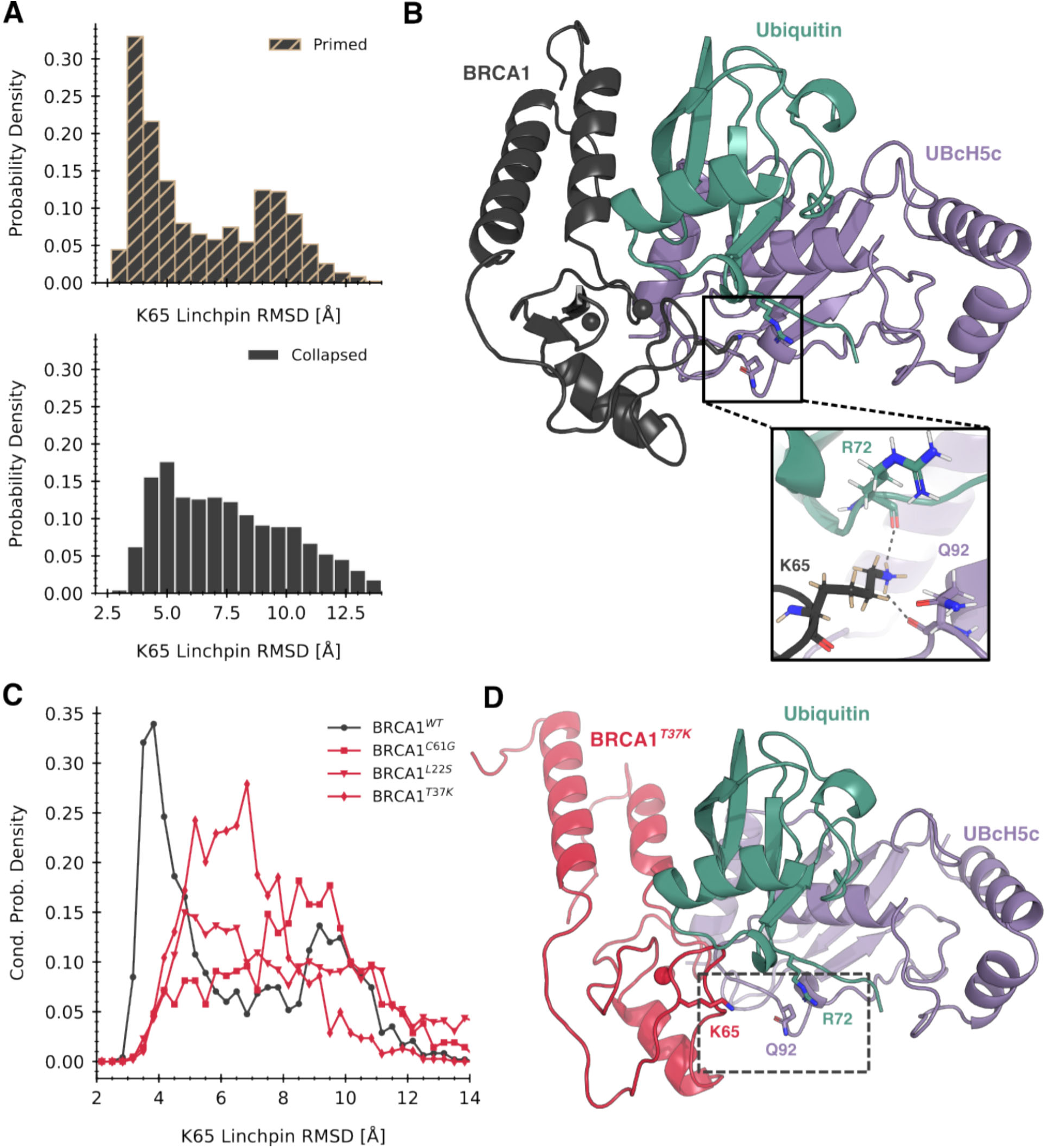
The likelihood of optimal linchpin geometry in monomeric states is significantly reduced by pathogenic mutations. A) Distribution of the RMSD of the K65 linchpin residue with respect to a structural model of the active state complex (part C) conditioned on primed and collapsed states. Primed states show a significant left shift in RMSD in the wild-type system. B) A structural model for the BRCA1 stabilizing the closed state of a UBcH5c conjugated to ubiquitin (BARD1 omitted for clarity). The inset shows the modeled hydrogen bonding network between the linchpin and residues on both UBcH5c and Ubiquitin. C) Comparison of conditional linchpin RMSD distributions between wild-type and pathogenic variants. All pathogenic variants show a pronounced right shift compared to wild-type, indicating they spend even less time in E2-binding competent conformations. D) A representative frame of the collapsed state of BRCA1^T37K^ aligned to the active complex. The linchpin and its hydrogen bonding partners are shown in spheres. The conformation of the E2 interface in a collapsed state results in a linchpin geometry far from the E2∼Ub interface, which makes formation of the hydrogen bonding network impossible.

We found that wild-type *BRCA1* has a much higher probability of proper linchpin priming in the primed states selected for by *BARD1*-binding. The conditional distribution of linchpin RMSD in the primed states exhibits significantly more mass at lower values compared to the collapsed states (Fig. 1a), indicating that primed states encompass conformations which are closer to the active state model (Fig. 4b). This finding highlights a direct mechanism for how *BARD1*-binding activates *BRCA1*. *BARD1*-binding induces the helix bundle into a primed state. Rotation of the helix bundle allosterically induces a conformational change in the *RING* domain, which shifts the K65 linchpin into a more optimal position to stabilize the E2∼Ub conjugate’s “closed” state. This in turn leads to higher rates of Ub transfer to the substrate protein.

Furthermore, we found that proper linchpin priming was less probable in primed states in the ensembles of all three pathogenic variants compared to primed states of wild-type (Fig. 4c). All three pathogenic mutations disrupt the shift towards optimal linchpin priming induced by stabilizing primed states of *BRCA1*, rendering binding with *BARD1* unproductive if it occurs. Suboptimal linchpin priming results in the relative position of K65 to be far from the E2/Ub interface (Fig. 4d). Therefore, the hydrogen bonding network which stabilizes the “closed” state of the E2∼Ub conjugate cannot form, reducing the rate of downstream Ub transfer. A reduction in the probability of linchpin priming is a common structural mechanism of pathogenesis for missense mutations in the N-terminal domain of *BRCA1*.

### Hyperactive mutations allosterically stabilize linchpin priming

Given that reduced linchpin priming in pathogenic mutants leads to a loss of function, we reasoned that enhanced linchpin priming would result in hyperactivation. Previously, Stewart et al. found that two mutations to *BRCA1* increase auto-ubiquitination activity above wild-type levels: L51W and K65R. Furthermore, they reported that the double mutant L51W/K65R was more active than either single mutant, and capable of restoring wild-type activity when introduced in conjunction with C61G^41^. We became interested in the *BRCA1*^L51W/K65R^ variant because of this reported rescue and conducted an additional ∼200 µs of simulation for it. Utilizing the same method as before, we analyzed the extent of linchpin priming in primed and collapsed states by comparing the distribution of linchpin RMSDs after alignment to the reference model.

As expected, we found that the hyperactivating mutation L51W/K65R has an even higher probability of adopting the primed linchpin than wild-type *BRCA1*. We found that the ensemble of the L51W/K65R variant exhibited enhanced linchpin priming independent of a collapsed or primed helix bundle (Fig 5a). The distribution of linchpin RMSD is left-shifted regardless of helix bundle state, leading to a net higher probability of linchpin priming. While the K65R mutation is known to drive higher catalytic activity due to stronger engagement with the E2/Ub interface, no explanation has been offered for the contribution of the L51W mutation. We determined that two key interactions between W51 and Zinc Finger II drive enhanced linchpin priming. First, substitution to tryptophan introduces additional hydrophobic bulk that contacts the sidechain of P59. This contact occurs more frequently in the L51W/K65R variant than it does in the wild-type *BRCA1* monomer or *BRCA1*/*BARD1* heterodimer (Fig. 5b). Second, the indole nitrogen in tryptophan is capable of making a hydrogen bond with the backbone of Q62 (Fig. 4c), which is impossible when a leucine is present. The combination of these two interactions stabilizes the structural states where the loop containing the linchpin residue is closer to the central α-helix, priming the linchpin.

**Figure 5.**
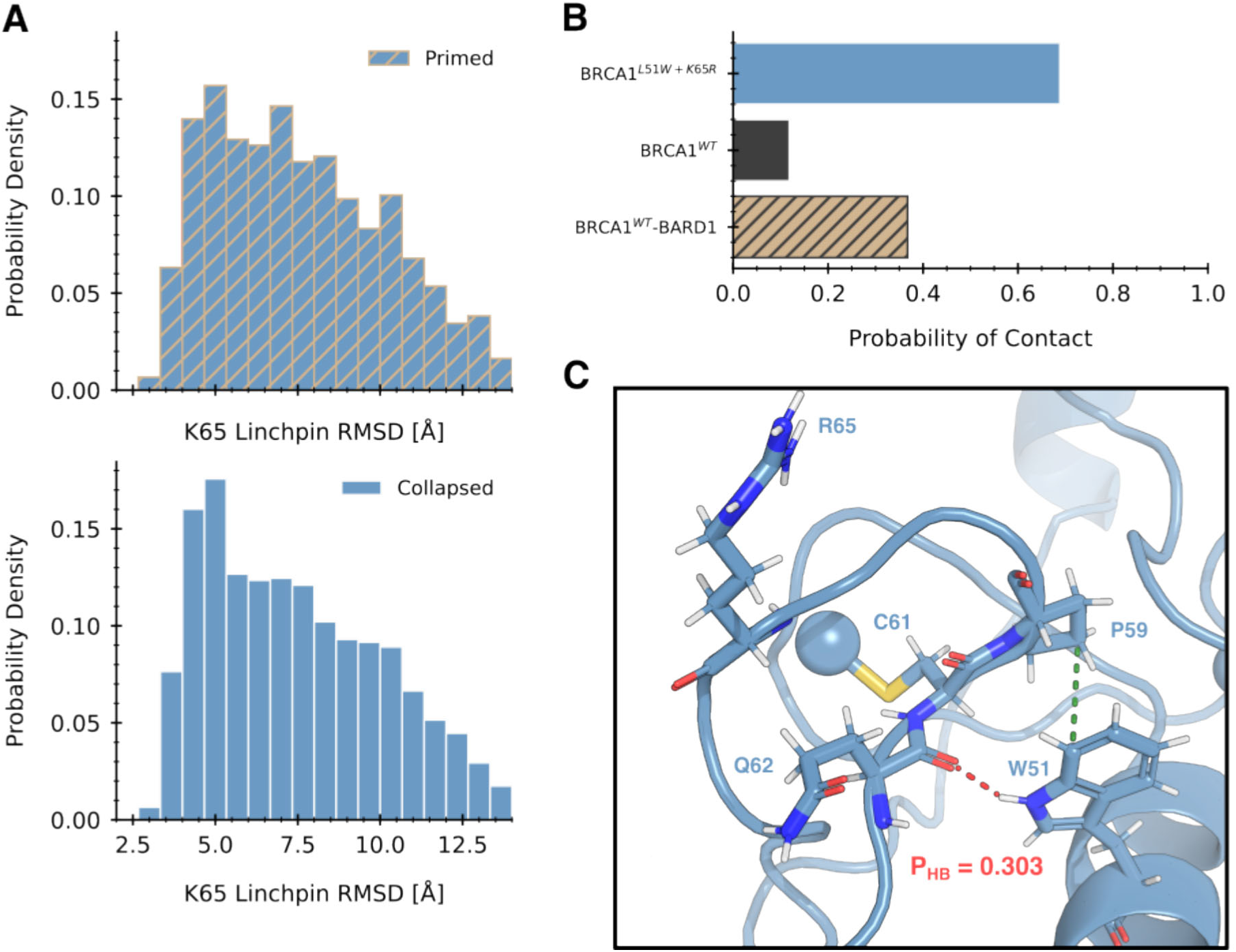
Hyperactivity of the L51W+K65R double mutant results from more linchpin priming in the monomeric state. A) Distributions of linchpin RMSD between primed and collapsed states of the L51W+K65R double mutant. Unlike wild-type (4a), there is almost no difference in the likelihood of linchpin priming between the two states. B) Probability of a contact between residue 51 and PRO 62. The L51W mutation allows a stronger hydrophobic interaction with PRO 62 (the green contact in the inset in panel C), stabilizing a zinc finger conformation amenable to priming. C) Example structure of an L51W-K65R double mutant aligned to the model of the quaternary complex. An additional hydrogen bond to the backbone oxygen of Q60 can be formed by the tryptophan mutation which is impossible in wild-type. This hydrogen bond is observed in approximately 30% of frames, lending additional stability to the primed state.

## Discussion/Conclusion

Understanding the molecular mechanism by which missense mutations lead to loss of function is a major advantage for the development of successful therapeutics. Developments in structural biology^43^, computational methods^44^, computing power^45^, and artificial intelligence^46^ have unlocked new ways of turning a mechanistic insight into a functionally active molecule. Gaining these insights, however, can be challenging if structure alone does not provide the necessary answers. Here, we utilize molecular dynamics simulations to unveil the structural mechanism for pathogenesis in the N-terminal domain of *BRCA1*.

The ensemble of the monomeric *BRCA1* has minimal overlap with heterodimeric *BRCA1* because of a collapse in the helix bundle. Our findings illustrate that as a monomer, *BRCA1* explores a conformational landscape which is both different and more varied compared to the heterodimer. The distinct monomer landscape arises from a twist in the helical bundle caused by the hydrophobic surfaces of the helices collapsing to mutually bury each other. These collapsed states are significantly more populated in the ensemble than the primed states, which are almost exclusively populated by the *BRCA1*/*BARD1* complex.

*BARD1*-binding to the helix bundle stabilizes a conformational change in the E2 interface of *BRCA1*. By deep learning features that report on structural elements of the protein, we identified a population shift in the *RING* domain conformation which is more likely when collapsed states are occupied. Our simulations revealed that the β-sheet in the *RING* domain is more likely to distort when the helix bundle twists into itself, repositioning the E2-interface.

We found that the extent of allosteric priming of the *BRCA1* linchpin residue at the E2 interface by *BARD1* distinguishes pathogenic and hyperactive variants from wild type. In primed states stabilized by *BARD1*, the extent of linchpin priming in wild-type *BRCA1* was significantly increased compared to collapsed states. Further comparison of collapsed-state linchpin positioning in wild-type and mutant ensembles to an active state model showed that all pathogenic mutations reduce the likelihood that the linchpin engages the E2/Ub interface. Additionally, we found that a hyperactive double mutation increases the likelihood of priming in either collapsed or primed states.

By clarifying the mechanism of pathogenesis, our work suggests a concrete strategy for the design of small molecules that directly restore the E3 ligase activity of N-terminal *BRCA1* mutants. A small molecule which either affects increased linchpin priming or mimics linchpin priming by stabilizing the E2∼Ub closed state at the interface with *BRCA1* is a potential avenue to achieve “structure correction.” The latter may be possible by the use of molecular glues^47^ at the heterotrimeric interface, while the former would require allosteric modulation of the E2 interface.

Application of the computational methods utilized in this work can generalize to any disease model which implicates missense mutations in a protein. Mutation-induced perturbations to protein dynamics have been demonstrated to be a targetable feature for precision drug discovery^48–51^. Although we have targeted this effort to *BRCA1*, owing to its importance in hereditary breast and ovarian cancer^10,52^, our approach serves as a blueprint for uncovering disease mechanisms related to protein variants. Future work may aim for the development of computational methods that increase the computational efficiency of this process. Lastly, we hope our contribution foments the development of effective treatments for *BRCA*-negative cancer patients.

## Methods

### Molecular Dynamics Simulations

Production molecular dynamics simulations were run with GROMACS^53^ on an in-house commodity cluster composed of NVIDIA RTX A5000, Quadro RTX 6000, and Tesla P100 GPUs, as well as CPU resources donated by Folding@Home^54^. All simulations were started from the NMR structure^19^ of the *BRCA1*/*BARD1* heterodimer reported by Brzovic et al. Mutant systems were constructed using the mutation wizard in PyMOL. Systems were prepared for simulation using AmberTools^55^. Each protein system was solvated with explicit water in a truncated octahedral box that extended 10 Å beyond the protein. Each system was neutralized with NaCl at a concentration of 150 mM. Systems were parameterized with the AMBER ff19SB^56^ forcefield, the OPC^57^ water model, and EZAFF^58^. OpenMM^59^ was used to minimize and equilibrate prepared systems. The steepest descent algorithm was to minimize each system until the maximum force fell below 10 kJ/(mol*nm). Then, each system was equilibrated, with heavy atoms restrained in place, with 1 ns of NVT and 1 ns of NPT simulation using the Berendsen^60^ thermostat and the Parrinello–Rahman barostat^61^ at 310 K. Production simulations were run in the NPT ensemble at 310K with the stochastic v-rescale^62^ thermostat and Parrinello-Rahman barostat. Simulations were initiated in rounds with adaptive sampling using the FAST algorithm^63^. Following that, simulations were seeded from representative frames of conformational states identified by clustering (see Data Analysis & Visualization).

### Structural Modeling of the BRCA1 Active State

A model for an active state *BARD1*/*BRCA1*/*UBcH5c*/*UB* complex was constructed in PyMOL to assess linchpin priming. A structure of the *RNF4*/*UbcH5a*/*Ubiquitin* complex^42^ was aligned to the *BRCA1*/*BARD1*/*UBcH5c* from PDB code 7LYB. Then, all available experimental structures of *BRCA1/BARD1* were aligned to 7LYB as well to find a model with the appropriate linchpin hydrogen bonding network. Model 2 from PDB code 1JM7 was ultimately used and replaced the heterodimer from PDB 7LYB. Sidechain rotamers were adjusted to fix any steric clashes.

### Data Analysis & Visualization

All data analysis was conducted using python, and all structures were visualized using PyMOL. Markov state models^32^ (MSMs) were constructed from simulation data for all systems. All phi, psi, chi-1 angles of *BRCA1* residues 16 to 85 were computed using mdtraj^64^ and used as input features. The dimensionality of this space was reduced to 10 dimensions using TICA^65^ (with a lag time of 10 ns) and then clustered using the implementation of the KMeans algorithm in deeptime^34^. Models were fit using row normalization of transition counts matrix. Model lag time for each system was selected using the implied timescales test. The number of centers was determined by cross validation^66^ with the rank-10 VAMP-2 score^67^. Computations were parallelized with jug^68^ where feasible. Once MSMs were finalized, a set of 30,000 samples was drawn from each system’s model. Each sample was generated by picking a conformational state according to the equilibrium distribution of the model, and then uniformly selecting a frame from all frames assigned to that state. All analysis characterizing ensembles was conducted on these representative samples. Root-mean-squared deviation, secondary structure (using the DSSP algorithm^35^), and solvent exposed surface area calculations were all done using implementations in mdtraj^64^. The linchpin RMSD was computed by first aligning a given frame to the reference model using alpha carbons in the secondary structure elements of the *RING* domain, then computing the displacement of the heavy atoms in lysine or arginine. A DiffNet^37^ was trained on the representative samples for wild-type monomer and heterodimer systems. The network was trained for 50 epochs on all *BRCA1* non-symmetric heavy atoms, with EM bounds of [0.2,0.8] and [0.6,0.8] for the monomer and heterodimer. Initial labels were 0.5 and 1.

## Acknowledgements

We thank the citizen scientists of the Folding@home community for their computing resources and support. In addition, we thank AMD for the donation of critical hardware and support resources from its HPC Fund that enabled many of the computations for this work. This work was supported by the Basser Center for BRCA at Penn Medicine and NIH grant R35GM152085. We additionally thank them for their insights and feedback on the manuscript.

## Author Contributions

AB performed all simulations and analyses, prepared figures, and drafted the manuscript. GRB proposed the initial project idea, gave feedback on project direction, provided edits to the manuscript, and acquired funding.

